# Effect of Aging on the Human Myometrium at Single-Cell Resolution

**DOI:** 10.1101/2023.07.03.547452

**Authors:** P Punzón-Jiménez, A Machado-Lopez, R Pérez-Moraga, J Llera-Oyola, D Grases, M Galvez-Viedma, M AlSibai, E Satorres, R Badenes, C Ferrer, E Porta-Pardo, B Roson, C Simón, A Mas

## Abstract

The myometrial dysfunction associated with aging can prompt complications during pregnancy and labor, causing a 7.8-fold increase in maternal mortality in women over 40. Using single-cell/single-nucleus RNA sequencing and spatial transcriptomics, we constructed a cellular atlas of the aging myometrium from 186,120 cells across twenty peri- and post-menopausal women. We identified 23 myometrial cell subpopulations, including novel contractile capillary, venous capillary, immune-modulated fibroblasts, and nervous system regulatory fibroblasts. Myometrial aging leads to fewer contractile capillary cells, a reduced level of ion channel expression in smooth muscle cells, and impaired gene expression in endothelial, smooth muscle, fibroblast, perivascular, and immune cells. We observed altered myometrial cell-to-cell communication as an aging hallmark associated with the loss of 25/229 signaling pathways, including those related to angiogenesis, tissue repair, contractility, immunity, and nervous system regulation. These insights may contribute to a better understanding of the complications faced by older women during pregnancy and labor.

## Introduction

The recent upward trend in human longevity provides significant challenges for healthcare systems worldwide, especially when considering women’s health^1^. According to the World Health Organization (https://www.who.int/news-room/fact-sheets/detail/ageing-and-health), 1 in 6 of the world’s population will be 60 years or over by 20230, with life expectancy for individuals in this age group expected to significantly increase^2^.

Aging impairs various physiological processes, including reproduction, pregnancy, and parturition^3^. Menopause, which typically occurs between the ages of 45 and 55, is caused by the irreversible ovarian demise that leads to the cessation of menses. Menopausal women experience various changes that significantly impact their everyday lives, such as hot flushes and night sweats^4^, vaginal dryness^5^, urinary incontinence^6^, weight gain^7^, and bone loss^8^.

Menopause was previously associated with the end of a woman’s reproductive lifespan^9^; however, the increased accessibility and availability of assisted reproductive techniques now support childbirth beyond this stage. These advances have largely bypassed infertility-associated difficulties related to advanced maternal age; however, the maternal mortality associated with the complications older women face during pregnancy and labor remains a significant problem. The 7.8-fold increase in maternal mortality in women aged >40 (107.9 deaths per 100,000 live births) compared to those under the age of 25 provides tangible proof of this problem^10, 11^.

The uterus is a contractile organ essential for reproduction, pregnancy, and labor. Various studies have linked dysfunction of the myometrium - the muscular outer layer of the uterus - to embryo implantation failure^12^, preterm birth^13^, labor dystocia ^14^, and uterine atony^15^, which all have significant implications for women’s health^16^. The thinning of the myometrium during menopause^17^ occurs alongside alterations to the vasculature^18^ fibrosis^19^, and reduced contractility^20^; however, we currently lack in-depth knowledge regarding the effects of age on the myometrium and how this impacts myometrial dysfunction.

Here, we performed single-cell/single-nucleus RNA sequencing (scRNA-seq/snRNA-seq) and spatial transcriptomics in the human myometrium during menopause to generate comprehensive cellular, molecular, and cell-to-cell communication (CCC) landscapes and understand the drivers of age-related myometrial dysfunction. We now report significant age-related alterations in cell type abundance and gene expression profiles in distinct cell types and validate these findings using spatial transcriptomics and CCC dynamics.

## Results

### Integrated Single-cell Atlas of the Human Myometrium during Menopause

To assess the cellular-level effects of aging on the myometrium, we examined myometrial tissue from the fundus, anterior and posterior wall of the uterus from 20 patients. Among these, 6 were obtained from perimenopausal women (≤54 years old), and 14 from postmenopausal women (>54 years old). Depending on tissue availability, biopsies were collected from the fundus and the anterior and posterior walls of the uterus.

Initially, we characterized the cellular heterogeneity of the entire myometrium by integrating data from uterine zone and menopause state. Our dataset comprised scRNA-seq data from 161,202 cells and snRNA-seq data from 24,918 nuclei, which provided an integrated single-cell cellular atlas that captures the diverse cellular landscape of the menopausal human myometrium (**Supplementary Fig. 1A**, **Supplementary Table 1**). After rigorous quality control filtering (see **Methods**) and batch correction (**Supplementary Fig. 1B**), we integrated data from the distinct uterine locations (anterior, posterior and fundus) and scRNA/snRNA-seq.

Cell clustering represented in the uniform manifold approximation and projection (UMAP) space and manual curation of canonical markers ^21^ prompted the identification of five major cell types: endothelial cells, fibroblasts, smooth muscle cells (SMCs), perivascular (PV) cells and immune cells, split into myeloid, lymphoid and mast cells (**Fig. 1A**). The remaining clusters represented cells of the epithelium, lymphatic endothelium (LEC), and peripheral nervous system (PNS) (**Fig. 1A**, **Supplementary Fig. 1C**).

**Figure 1.**
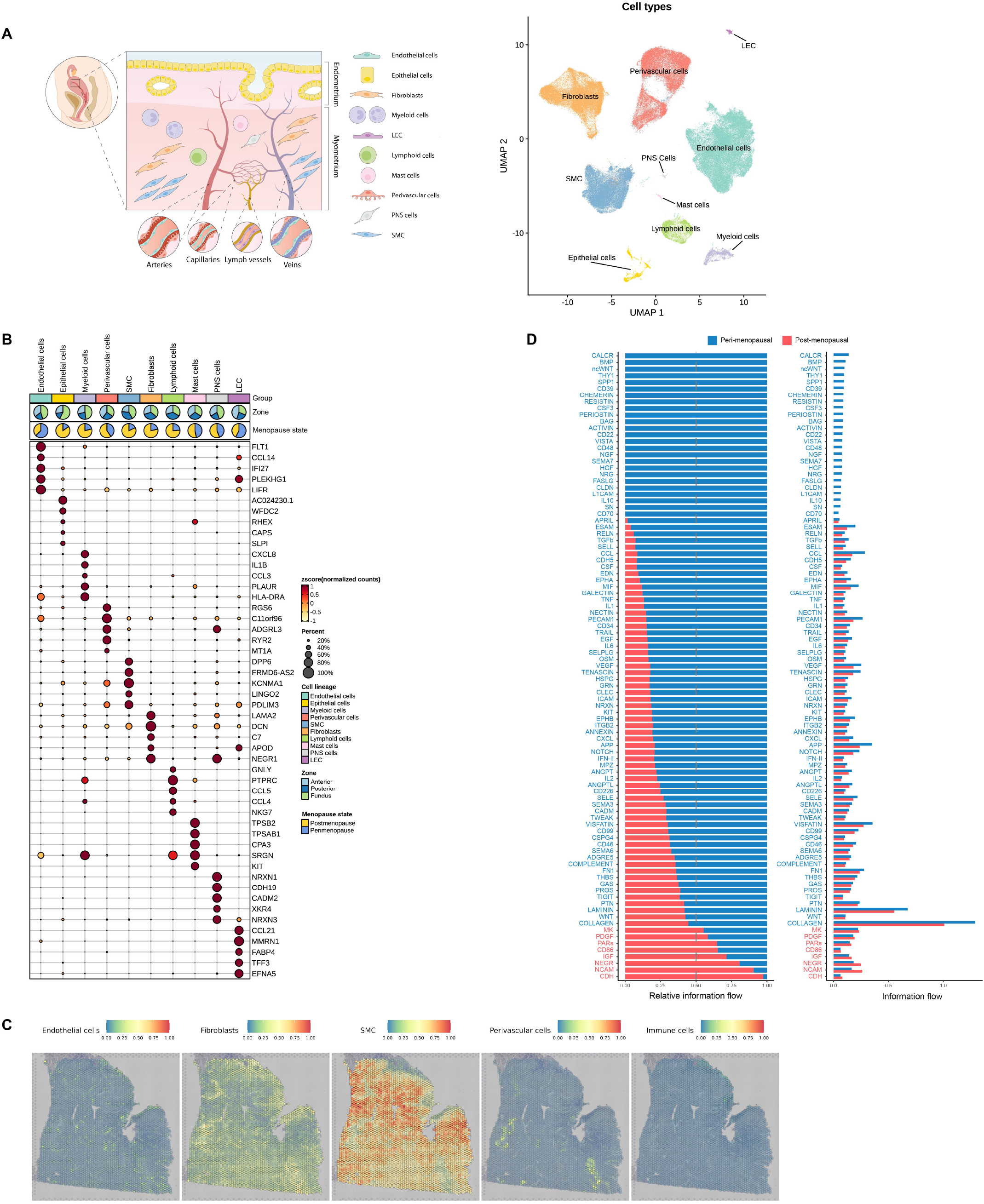
Integrated Single-Cell Atlas of the Human Myometrium. **A)** Schematic representation of the human uterus and the identified cell types within the myometrium (left). Visualization of uniform manifold approximation and projection (UMAP) showing an integrative clustering of high-quality cells and nuclei from human myometria (peri-menopausal, n=6; post-menopausal, n=14) (right). **B)** Dotplot indicating the relative expression of each identified cell type’s top five discriminatory genes (Color indicates average expression, while dot size represents the percentage of cells expressing specific genes). Pie charts on top illustrate the proportion of cells according to menopausal state (peri-menopause in blue, post-menopause in yellow) and myometrial zone (light blue anterior region, dark blue posterior region and green fundus) **C**) Panels displaying the spatial location of endothelial cells, fibroblasts, SMCs, PV cells, and immune cells in the myometrium. **D)** Relative and absolute flows of differentially active signaling pathways during myometrial aging (comparing peri- and post-menopause myometria). Abbreviations: LECs: lymphatic endothelial cells; PNS: peripheral nervous system; SMCs: smooth muscle cells.

The endothelial cell population was characterized by the expression of *FLT1, CCL14, IFI27, PLEKHG1, LIFR* (**Fig. 1B**) and canonical genes such as *VWF* and *ENG* (**Supplementary Fig. 2A**), which participate in the maintenance of hemostasis through vasculogenesis, blood vessel morphogenesis, and angiogenesis. The fibroblast population specifically expressed *LAMA2, DCN, C7, APOD, NEGR1* (**Fig. 1B**), and *LUM* (**Supplementary Fig. 2A**), which function in collagen organization, immune response, and cell adhesion. The SMC population overexpressed *DDP6, FRMD6-AS2, KCNMA1, LINGO2,* and *PDLIM3* (**Fig. 1B**), which participate, together with canonical genes such as *ACTA2* and *DES* (**Supplementary Fig. 2A**), in muscle structure development and contractility-associated functions, including ion transport, depolarization, and action potential through calcium and potassium channels. The PV cell population expressed increased levels of *RGS6, c11orf96, ADGRL3, RYR2,* and *MT1A* (**Fig. 1B**), which, alongside *MYH11* and *ACTA2* (**Supplementary Fig. 2A**), regulate G protein-coupled receptor signaling cascades and calcium channel activity. The immune cell population represented a heterogeneous group that overexpressed canonical markers such as *CD3E, TRBC2, AIF1, CYBB, TPSB2,* or *TPSAB1* (**Fig. 1B**, **Supplementary Fig. 2A**). Myeloid cells were characterized by *CXCL8, IL1B, CCL3, PLAUR*, and *HLA-DRA* expression, which associates with inflammatory stimuli and responses by attracting neutrophils and basophils. Lymphoid cells expressed *GNLY, PTPRC, CCL5, CCL4,* and *NKG7*, which relate to cytolytic T-lymphocytes and NK cells and represent essential T- and B-cell antigen receptor signaling regulators. Meanwhile, mast cells expressed *CPA3, SRGN,* and *KIT*, which play roles in innate immunity.

Subsequently, we systematically mapped the spatial distribution of the main cell populations identified using sc/snRNA-seq within the tissue architecture of the myometrium (**Fig. 1C**). Our findings revealed the wide distribution of fibroblasts and SMCs, while endothelial and PV cells displayed a more restricted distribution, confined to regions associated with blood vessels. We also confirmed the presence of a small subset of epithelial cells correlating with remaining endometrium by histological examination (**Supplementary Fig. 2B**). These results provided a rationale for excluding remaining endometrium from further analyses.

Lastly, given the critical role of CCC in maintaining tissue homeostasis, we examined interactions between distinct myometrial cell types and explored potential age-related disruptions. Specifically, we inferred all intercellular communications by analyzing the expression of ligand-receptor pairs using CellChat^22^. Interestingly, CCC networks revealed the complete loss of signaling pathways related to contractility, angiogenesis, tissue repair, nervous system regulation, and anti-inflammatory processes in the post- menopausal myometrium, suggesting that aging leads to myometrial dysfunction (**Fig. 1D**).

These data demonstrate the cellular heterogeneity of the menopausal myometrium, which comprises ten distinct cell types, and highlight significant alterations to CCC linked to aging.

As the next step, we explored the potential impact of aging on individual cell types within the myometrium. We focused on each major cell type and carefully delineated the cell subtypes by curating gene markers and then conducted comparative analyses to discern alterations in abundance, gene expression profile, CCC, and spatial distribution of each cell subtype between the peri- and post-menopausal myometria as our model of aging.

### Endothelial dysfunction associated with the aging myometrium

We assessed the impact of aging on myometrial endothelial cells through an analysis of four main subpopulations – arterial, contractile capillary, venous capillary, and venous cells (**Fig. 2A**). Arterial cells expressed *FN1, IGFBP3, NEBL,* and *ARL15*; contractile capillary cells expressed *TPM2, ACTA2, TAGLN, MYH11*, and *MYL9*; venous capillary cells expressed *SELE, CCL2, TM4SF1*, and *ADAMTS9*; and venous cells expressed *TXNIP, NOSTRIN, LIFR, ACKR1*, and *CCL14* (**Supplementary Table 3**). Differential abundance analysis (**Fig. 2B**), found significant changes within the endothelial cell populations associated with myometrial aging, as depicted in the neighborhood graph and beeswarm plot. Notably, we discovered a lower abundance of contractile capillary cells in post-menopause myometria (**Fig. 2C**), while differential gene expression analysis revealed deregulated gene expression profiles in post-menopause vs. peri-menopause myometria (**Supplementary Table 4**). Specifically, arterial cells overexpressed *CCN1,* venous, venous capillary, and contractile capillary cells overexpressed *NFATC2*, while arterial, venous, and venous capillary cells overexpressed *ANO2* in post-menopausal myometria (**Fig. 2D**). We also observed the downregulation of *TAGLN* in venous and venous capillary cells, *MYL9* and *MYH11* in venous capillary cells, and *ACTG2* and *FLNA* in contractile capillaries compared to peri-menopause samples (**Fig. 2D**). Furthermore, using spatial transcriptomics and a refined spatial map approach for improved resolution, we confirmed the significant reduction in the contractile capillary subpopulation of endothelial cells within the post-menopausal myometrium (n=5) as compared to the peri-menopausal myometrium (n=3). (Fig. 2E and F).

**Figure 2.**
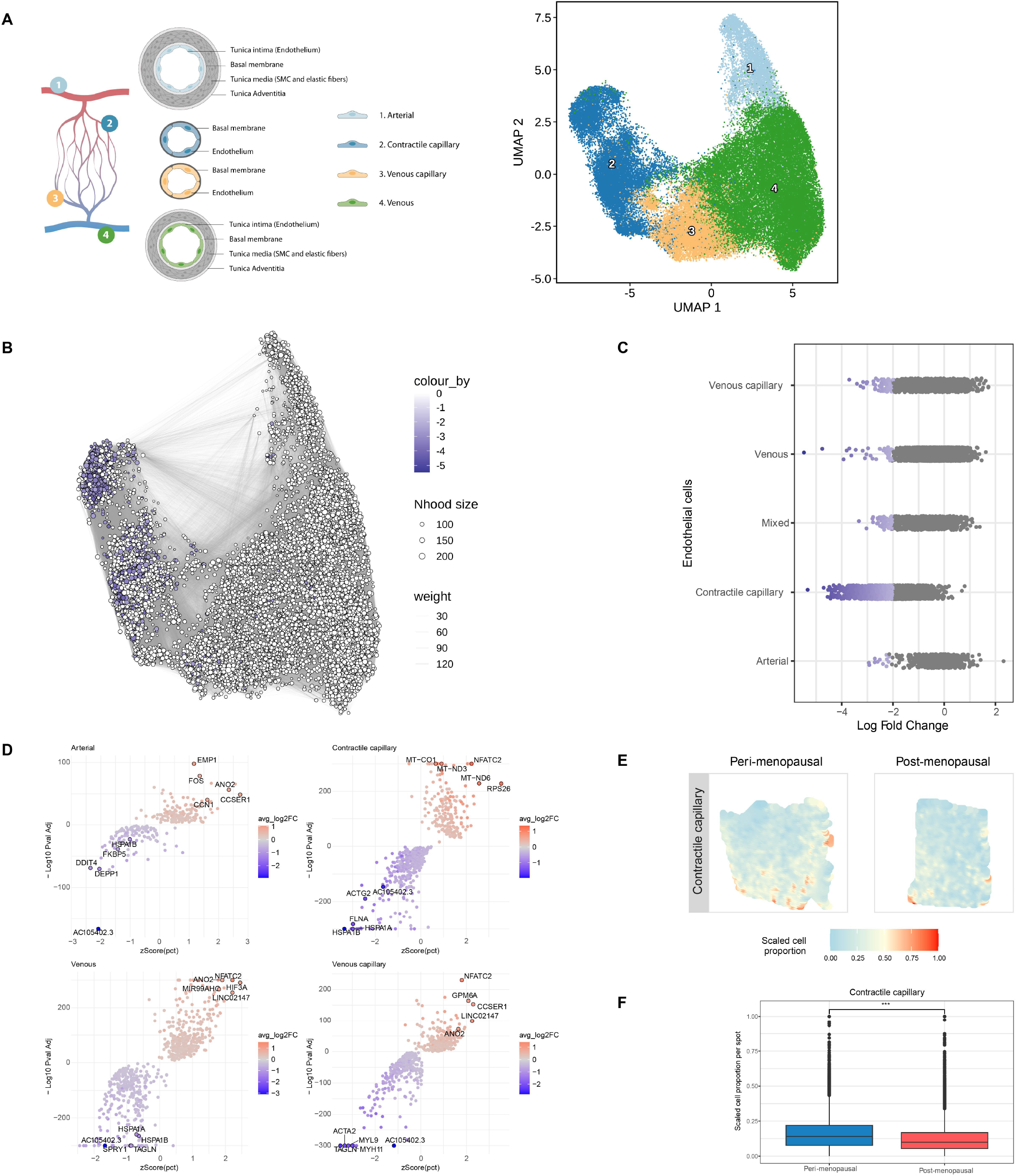
Endothelial Dysfunction in the Aging Myometrium. **A)** Schematic representation of the endothelial cells (left) and UMAP visualization of the principle endothelial subpopulations (right) in the myometrium. **B)** Neighborhood graph highlighting the differential abundance of endothelial cells in the aging myometrium. Dot size represents neighborhoods, while edges depict the number of cells shared between neighborhoods. Neighborhoods colored in blue represent those with a significant decrease in cell abundance during myometrial aging. **C**) Beeswarm plot of differential cell abundance by cell type. X-axis represents the log-fold change in abundance during myometrial aging. Each dot represents a neighborhood; neighborhoods colored in blue represent those with a significant decrease in cell abundance in post-menopause myometrial samples. **D**) Volcano plots representing age-related differentially expressed genes in each endothelial cell subpopulation. Positive LogFC indicates overexpression in the post-menopausal myometrium, whereas negative LogFC indicates overexpression in the peri-menopausal myometrium. **E)** Representative refined spatial maps of contractile capillary cells in peri-(left) and post-menopausal (right) myometrium. Color represents the scaled proportion of this cell type in each location (red indicates the highest proportion of contractile capillary cells in that tissue). **F)** Boxplot of the scaled cell proportions of contractile capillary per spot split in the post- and peri-menopausal myometrium (***= p value<0.001). Abbreviations: SMC: smooth muscle cell; Nhood size: Neighborhood size.

These findings suggest that aging is associated with an impairment in the myometrial endothelial cell population; specifically, we observed both a reduced number of cells and the reduced expression of contractility-associated genes such as *FLNA* and *ACTG2*.

### Age-related transcriptomic alterations within the myometrial fibroblast population

We identified eight distinct fibroblast subpopulations within the myometria based on their transcriptomic footprint (**Fig. 3A**). Analysis of gene expression for each subpopulation (**Fig. 3B, Supplementary Table 5**) revealed that classic fibroblasts expressed *RARRES2, SELENOP, RPS10, MPG,* or *APOD*, while the expression of *CCL2, GPRC5A, CYTOR, or IL1R1* characterized immune-modulated fibroblasts. Intermediate fibroblasts specifically expressed *NRP1,* while myofibroblasts expressed *MYH11, DPP6,* and *KNMA1*. Nervous system regulatory fibroblasts expressed *APOD, PI16, EBF2, CDH19*, and genes associated with nervous system regulation (such as *NLGN1* or *NRP2*). Stressed fibroblasts expressed genes involved in stress-response and DNA damage, such as *HSPA1A, HSPA1B, HSPA8, DNAJB1,* and *GADD45B*. Additionally, we labeled two further subclusters of “universal” fibroblasts based on the expression of *COL15A1* and *PI16*^23^.

**Figure 3.**
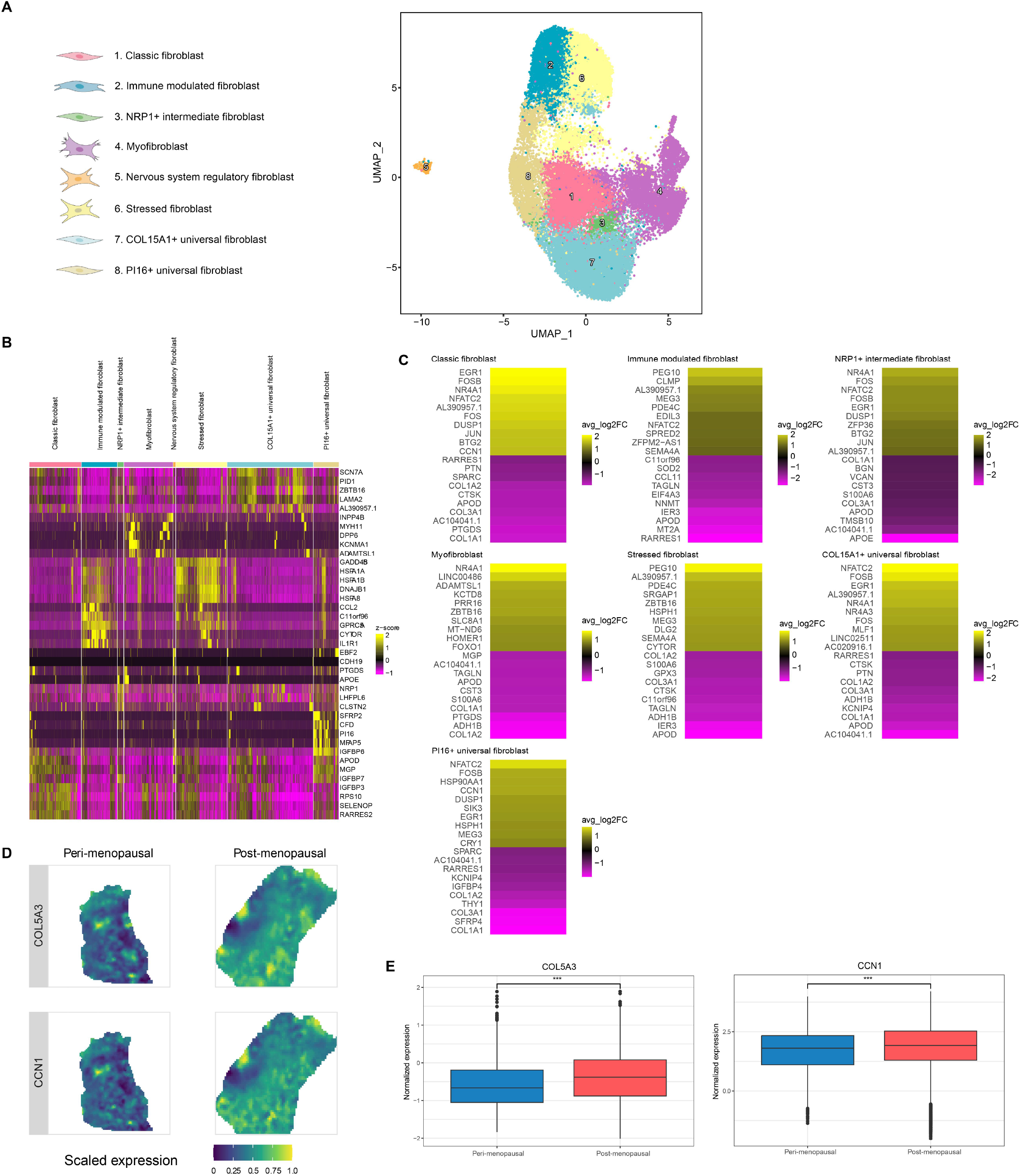
Age-Related Changes in the Myometrial Fibroblast Population. **A)** UMAP visualization displaying the fibroblast subpopulations in the myometrium. **B)** Heatmap showing the relative expression of the top five discriminatory genes in each subpopulation. **C)** Heatmap showing the genes with the top ten highest and lowest LogFC values in the post- vs. peri-menopause myometrium for each subpopulation. Positive LogFC indicates overexpression in the post-menopausal myometrium, whereas negative LogFC indicates overexpression in the peri-menopausal myometrium. **D)** Representative refined spatial maps of the expression of *COL5A3* and *CCN1* in peri-(left) and post-menopausal (right) myometrium. Color indicates the expression levels in each spot (yellow indicates the highest expression of each gene in the tissue). **E)** Boxplot of the spatial expression of the collagen *COL5A3* (upper) and fibroblast senesce marker *CCN1* (lower) genes in the peri- and post-menopausal myometrium (***= p value<0.001).

While we did not observe any age-related alterations to cell type abundance in the fibroblastic compartment during myometrial aging (data not shown), an analysis of the top ten most overexpressed genes in peri- (negative LogFC; purple) and post-menopause (positive LogFC; yellow) revealed an increase in aging-associated genes (e.g., *FOS* and *FOSB*) and immune system regulation genes (e.g., *NFATC2*) in universal PI16+ and COL15A1+, classic, and intermediate NRP1+ fibroblasts in the post-menopausal myometria (**Fig. 3C & Supplementary Table 6**). Furthermore, except for the nervous system regulatory fibroblasts, the remaining fibroblasts displayed a general downregulation in the expression of genes encoding proteins that participate in the formation and maintenance of collagen homeostasis (e.g., *APOD, COL1A1, COL1A2*, and *COL3A1*) during myometrial aging (**Fig. 3C & Supplementary Table 6**). Of note, the only differential expression observed in the nervous system regulatory fibroblasts was related to the *PABPC1* gene **(Supplementary Table 6**).

Spatial transcriptomics revealed similar cell densities and distributions in both peri- and post-menopausal myometrial and confirmed the increased expression of collagen (*COL5A3)* and senescence (*CCN1)* markers such during aging (**Fig. 3D and E**).

Overall, we did not observe changes to the abundance of specific fibroblast subpopulations associated during myometrial aging; however, gene expression analysis suggested significant changes in myofibroblast, universal (*COL15A1, PI16*), classic, immune modulated, stressed and NRP1+ intermediate fibroblasts.

### More Abundant but Less Contractile and Ion-conductive Smooth Muscle Cells Characterize Myometrial Aging

We identified four SMC subtypes - canonical, contractile, stimuli-response, and stressed – in the myometrium (**Fig. 4A**). The canonical SMC subtype expressed classical SMC markers, such as *ACTG2, MYL9*, *TPM2*, *TCEAL4, LGALS1, RPL30, S100A6*, and *CD81* (**Supplementary Table 7**). Contractile SMCs expressed contractility-associated genes (e.g., *CARMN, KANSL1, PBX1, LBD2, FRMD6-AS2*) and ion channel genes (e.g., *KCNMA1, CACNA1C,* and *CACNA1D*). Stimuli-response SMCs expressed *LMCD1*, *HRH1*, *MAFF*, *SAMD4A,* and *HOMER1*, which participate in inflammatory responses. Finally, stressed SMC expressed stress-associated genes (e.g., *JUN, JUNB*, *FOSB*, *ZFP36,* and *GADD45B*).

**Figure 4.**
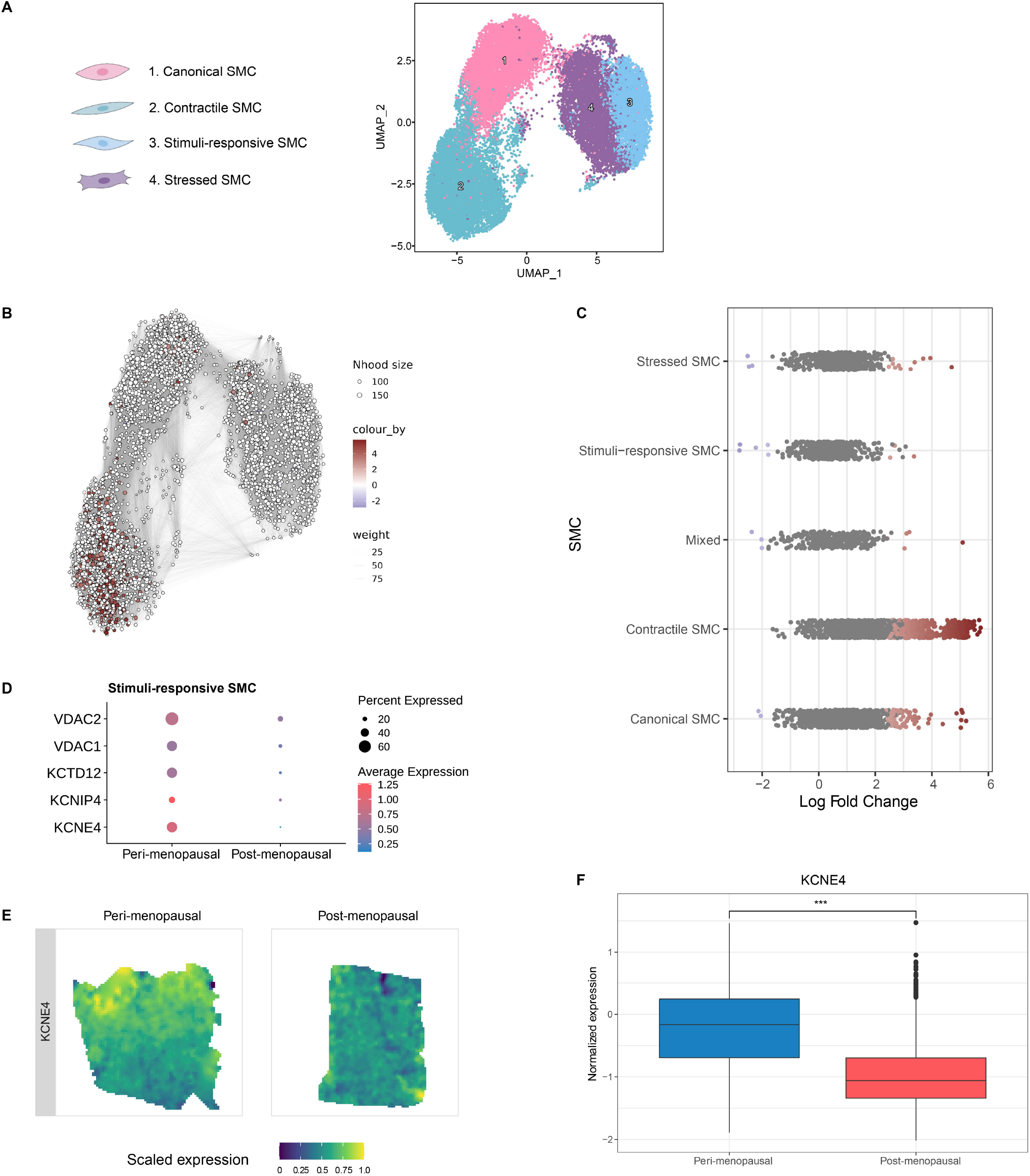
Increased Abundance but Reduced Functionality of SMCs Accompanies Myometrial Aging. **A)** UMAP visualization of the four distinct SMC subpopulations. **B)** Neighborhood graph highlighting the differential abundance of SMCs in the aging myometrium. Neighborhoods colored in dark red represent those with significantly increased abundance in the post-menopause myometrium. **C**) Beeswarm plot of differential SMC abundance by cell type. **D**) Dot plots representing the differential gene expression of voltage channel encoding genes during myometrial aging in stimuli-response SMCs. Dot size indicates the percentage of stimuli-response SMC that express the gene, while color indicates the average expression **E)** Representative refined spatial maps of the expression of potassium voltage-gated channel gene *KCNE4* in peri-(left) and post-menopausal (right) myometrium. Color indicates expression levelsin each spot (yellow indicates the highest expression of each gene in the tissue) **F)** Boxplot of the spatial expression of the KCNE4 gene in peri- and post-menopausal myometria. Abbreviations: SMC: smooth muscle cells; Nhood size: Neighborhood size.

Comparing alterations in the relative abundance of SMC subpopulations and their transcriptomic profiles during myometrial aging revealed a significant increase in the abundance of contractile and canonical SMCs (**Fig. 4B and C**). and the reduced expression of contractility-associated genes (e.g., *SMTN*, *ACTG2, MYL6, MYL9, TPM2,* or *DES*) in all SMCs during myometrial aging (**Supplementary Fig. 3A & Supplementary Table 8**). Interestingly, we found significant downregulation in the expression of K+ voltage channel genes (*VDAC1, VDAC2, KCTD12, KCNIP4, KCNE4*) in post-menopausal myometrial samples (**Fig. 4D**). Spatial transcriptomics analysis confirmed previous findings by demonstrating an increase in the density of SMCs in post-menopausal myometrial samples (**Supplementary Fig. 3B**), but the reduced expression of contractility marker genes such as *SMTN* (**Supplementary Fig. 3C**) and voltage channels such as *KCNE4* (**Fig. 4E and F**). Our results suggest that a greater abundance of SMCs with lower contractile and ion-conductive activities accompanies myometrial aging.

### Age-induced inflammation and DNA damage markers in myometrial perivascular cells

Analysis of PV cells in myometrium provided evidence of seven subtypes, four pericyte and three vascular smooth muscle cell (VSMC) populations (**Fig. 5A, Supplementary Table 9**). We identified the pericyte subtypes based on the expression of canonical markers *RGS5* and *PDGFRB*^24, 25^. The RGS5^+^PDGFRB^+^ subtype expressed *PDZD2* and *DLC1*; the RGS5^+^PDGFRB^−^ subtype expressed *CCL2, ADAMTS4, EMP1,* and *STEAP4*; and the RGS5^−^PDGFR^+^ subtype expressed *STEAP4, APOE, ADH1B,* and *ARHGAP15*. The RGS5^−^ PDGFR^−^ subtype (“contractile pericytes”) presented a partial loss in their canonical markers but a gain in the expression of contractility markers such as *DES* and *ACTG2.* The VSMC subtypes included contractile VSMCs expressing *ACTA2*, *ELN, CARMN,* and *KCNMA1*; damage-responsive VSMCs expressing stress response- and DNA repair-associated genes (*ATF3, FOS, JUN, IER2, KLF2)*; and stimuli-responsive VSMCs expressing genes related to the transmission of muscular and neural impulses (*ATP1B3* and *TNC*), arterial damage (*PVT1*), response to systemic inflammation (*CCL2)* and cell-cycle and transcription control (*CCNH* and *ZNF331)*.

**Figure 5.**
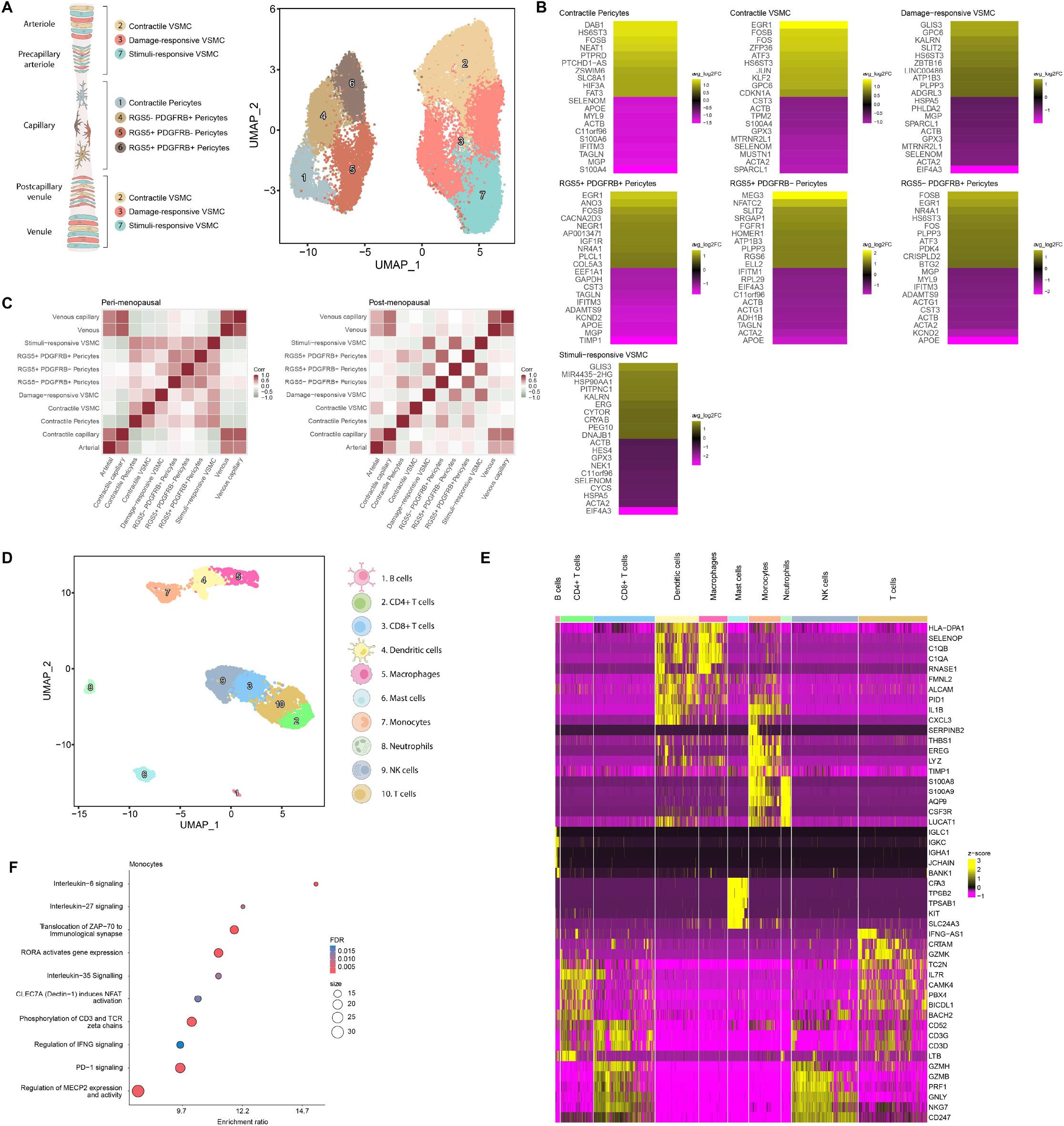
Impairments in the Myometrial Perivascular and Immune Cell Populations Associated with Myometrial Aging. **A)** (Left) Schematic representation of PV subtypes surrounding the myometrial microvasculature from the arteriole to the venule: arterioles and venules are surrounded by a single layer of contractile VSMCs, while pericytes (characterized by a stellate shape) are usually found surrounding smaller and transitional vessels such as precapillary arterioles and postcapillary venules. Contractile pericytes provide support in capillaries. (Right) UMAP visualization displaying the PV subpopulations. **B)** Heatmap showing the genes with the top ten highest and lowest LogFC values in the post- vs. peri-menopause myometrium for each PV subpopulation. Positive LogFC indicates overexpression in the post-menopausal myometrium, whereas negative LogFC indicates overexpression in the peri-menopausal myometrium. **C)** Correlation of the spatial distribution of endothelial and PV cell populations in the peri- (left) and post-menopausal (right) myometrium, with color scaled by correlation value (darker red indicates a higher correlation between the location of the indicated cell types). **D)** UMAP visualization displaying the immune cell sub-clusters present in the myometrium. **E)** Heatmap showing the relative expression of the top five discriminatory genes in each subpopulation. **F)** Dotplot showing the representative biological processes and pathways affected in monocytes during myometrial aging based on differential gene expression. Significant over-representation of biological processes and pathways (color intensity) shown by each gene set (dot size) from peri- and post-menopausal monocyte cells. Abbreviations: VSMC: vascular smooth muscle cells; NK: natural killer; FDR: false discovery rate.

Analysis of the top ten DE genes with the highest and lowest expression values in PV cells revealed that myometrial aging is associated with the upregulated expression of genes related to inflammation, immune response, and DNA damage but the downregulated expression of contractility genes associated with Ca2+/K+-related signaling, and vascular health-related genes (*MGP, TAGLN, ACTA2, ACTB, ACTG1, APOE, or KCND2*) (**Fig. 5B & Supplementary Table 10**).

We next created correlation plots for a more comprehensive spatial transcriptomic analysis to validate the co-localization of the PV and endothelial compartments, which revealed a decreased spatial co-localization of endothelial and PV cells during myometrial aging (**Fig. 5C**) and suggested the existence of an age-related impediment in the ability of PV cells to support endothelial cells.

Our findings indicate significant changes within the PV compartment of the aging myometria, which prompts a decline in the PV cells that support the endothelial compartment. Consequently, this compromise in PV support may adversely affect the optimal function of the endothelium.

### Immune cell cartography of the aging myometrium highlights monocytic dysfunction

Finally, we analyzed ten different subpopulations of immune cells (including myeloid, lymphoid, and mast cells) in the aging myometrium (**Fig. 5D**). Myeloid cells encompassed four subclusters (neutrophils, monocytes, dendritic cells, and macrophages) while lymphoid cells encompassed five subclusters (B cells, undifferentiated T cells, CD4+ T cells, CD8+ T cells, and NK cells) (**Fig. 5E, Supplementary Table 11**).

Although we did not observe significant variations in the abundance of specific cell subpopulations during myometrial aging (data not shown), differential gene expression analysis revealed significant differences observed mainly in the monocyte subpopulation (**Supplementary Table 12**). Monocytes overexpressed genes implicated in inflammatory processes and that facilitate immune responses through monocyte activation, adhesion, and migration (e.g., *PBX1, BNC2, EGR1, RORA*, and *DCN*). An overrepresentation analysis performed on monocytes revealed the enrichment of pathways associated with interleukin signaling, T-cell response modulation, and macrophage activation (**Fig. 5F**).

Our results suggest that myometrial aging affects monocytes gene expression that are responsible of maintaining the innate immune response by regulating T-cell activity and macrophage activation through cytokine signaling.

### Altered Cell-to-Cell Communication as a Hallmark of Myometrial Aging

Given the critical role of CCC in physiological tissue function, we next examined the impact of aging on CCC within the myometrium. In general, the post-menopausal myometrium prompted the appearance of less intricate signaling networks compared to the peri-menopausal myometrium (**Fig 6A, C, E, F, G, and I)** as well as shift in ligand-receptor pair interactions (**Fig 6B, D, H and J)**, which suggests that reduced CCC accompanies myometrial aging.

**Figure 6.**
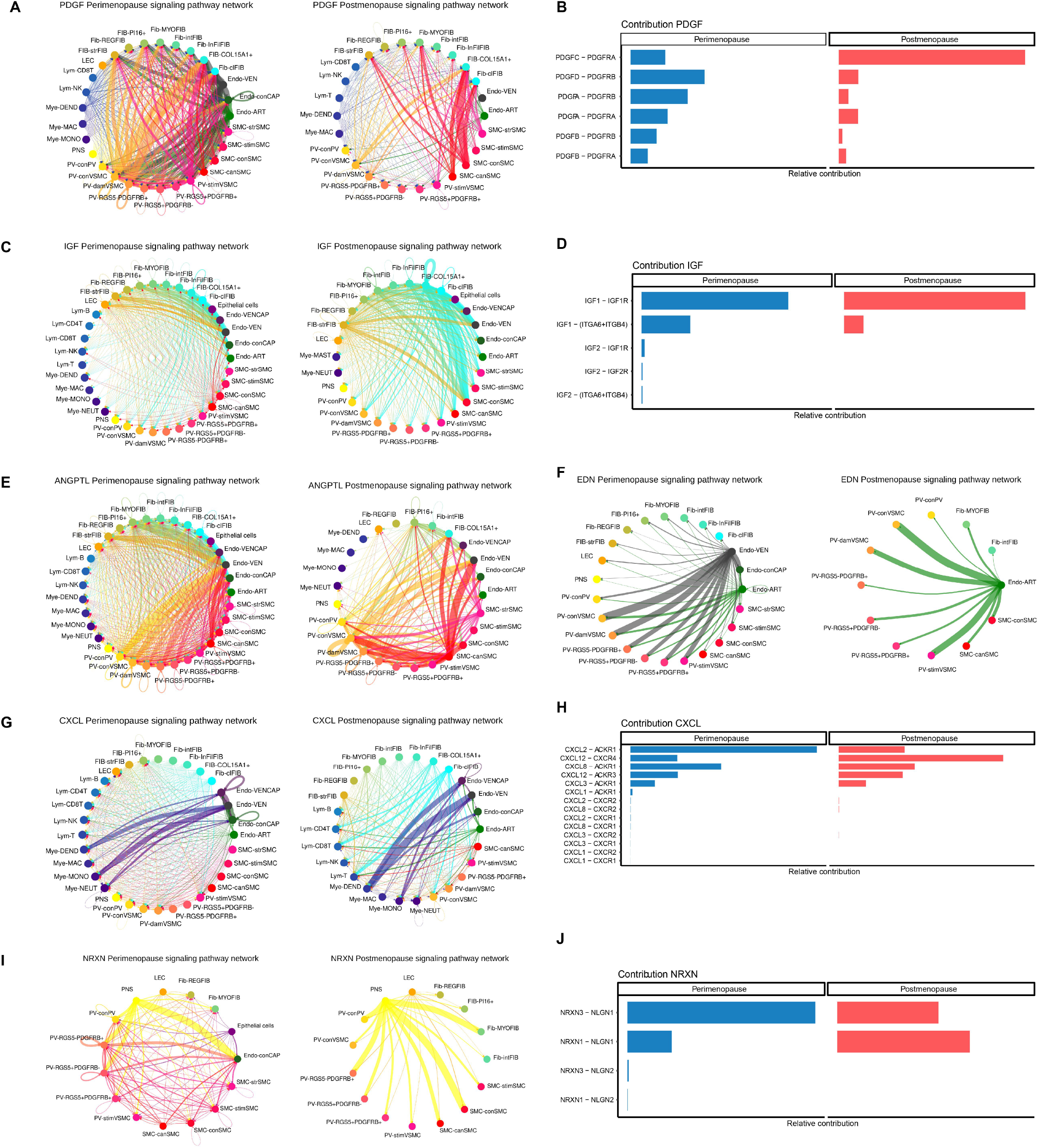
Age-Related Changes in Myometrial Cell-to-Cell Communication. Arrows between cell types are colored according to the cell type emitting the signal. The relative thickness of each line depicts the expression-based strength of the interaction between cell types. CCC chord plots for **A**) PDGF, **C**) IGF, **E**) ANGPTL, **F**) EDN, and **G**) CXCL signaling pathways in the peri-menopausal myometrium (left) and post-menopausal myometrium (right). The relative contribution of ligand-receptor pairs to CCC in the B) PDGF, D) IGF, H) CXCL, and J) NRXN signaling pathways in the peri- (left) and post-menopausal myometrium (right). Abbreviations: PDGF: platelet-derived growth factor; IGF: insulin growth factor; ANGPTL: angiopoietin-like; EDN: endothelin; CXCL: C-X-C Motif Chemokine Ligand; NRXN: nerve transmission-associated neurexin; Fib: fibroblasts; Endo: endothelial; SMC: smooth muscle cells; VSMC: vascular smooth muscle cells; PV: perivascular; LEC: lymphatic endothelium; Mye: myeloid; Lym: lymphoid; VEN: venous; ART: arterial; str: stressed; stim: stimuli-response; con: contractile; can: canonical; dam: damage-response; MAC: macrophages; DEND: dendritic; NK: natural Killer; MONO: monocytes; NEUT: neutrophils; REG: nervous system regulatory fibroblast; MYO: myofibroblast; int: NRP1 intermediate; Inf: immune modulated fibroblast; Cl: classic.

We identified 204 pathways were identified in both peri and post-menopausal myometria and highlighted potentially relevant pathways in processes comprising myometrial function during aging. These include as detailed hereafter, the fibrosis-associated platelet-derived growth factor (PDGF) and insulin growth factor (IGF) pathways, the angiogenesis-associated angiopoietin-like (ANGPTL) and endothelin (EDN) pathways, the inflammation-associated C-X-C Motif Chemokine Ligand (CXCL) pathway, and the neurexin (NRXN) pathway which plays a crucial role in impulse nerve transmission (**Supplementary Fig. 4A and B**).

In the aging myometrium, the PDGF signaling pathway suffered a notable reduction in the outgoing signals from the PVs and SMCs (**Fig. 6A**), and a shift in the relative contribution of specific ligand-receptor pairs (**Fig. 6B**). Interestingly, a shift from a more diverse distribution of ligand-receptor pairs and the contribution of the angiogenesis-related PDGFD-PDGFRB axis to a less diverse distribution of ligand-receptor pairs and the contribution of the fibrosis- and inflammation-associated PDGFC-PDGFR axis accompanied myometrial aging.

Analysis of the IGF pathway revealed an age-related increase in signals emanating from fibroblasts and SMCs (**Fig. 6C**). While aging did not induce widespread alterations in ligand-receptor pair interactions, we did observe the loss of IGF2-associated interactions and a subtle reduction of IGF1-IGF1R interactions with aging (**Fig. 6D**). Age-related alterations to the ANGPTL pathway primarily affected PV cells, endothelium, and SMC subtypes, with their decline suggesting a potential reduction of blood vessel sprouting (**Fig. 6E**), while a shift from EDN signaling in the arterial and venous endothelium to only the arterial endothelium accompanied the aging process (**Fig. 6F**).

We encountered notable alterations to the immune system-associated CXCL signaling pathway (**Fig. 6G**); we observed a shift from extensive communication between various cell types (considerably strong between myeloid cells and endothelial subtypes) to more robust communication of fibroblasts with immune and myeloid cells with endothelial subtypes during myometrial aging (**Fig. 6G**). We also observed a shift from the activity of the CXCL2-ACKR1 ligand-receptor pair in peri-menopausal myometria to CXCL12-CXCR4 in post-menopausal myometria. Both receptor pairs are associated with inflammation and immune activation (**Fig. 6H**).

Lastly, analysis of the impulse nerve transmission-associated NRXN signaling pathway suggested a shift to fewer but stronger interactions, particularly in signals transmitted from the PNS to PV, SMC, and fibroblast subpopulations during myometrial aging (**Fig. 6I**); Moreover, we identified a shift between interactions of the NRXN3-NLGN1 and NRXN1-NLGN1 ligand-pairs, which decreased and increased during aging, respectively (**Fig. 6J**).

Overall, CCC data reveals that a generalized decline in signaling complexity accompanies myometrial aging, which may prompt detrimental processes such as increased fibrosis, inflammation, impaired angiogenesis, and reduced responsiveness to stimuli.

### The Loss of Signaling Pathways Accompanies Myometrial Aging

The analysis of CCC revealed that aging associated with the loss of incoming and outgoing interactions from 25 signaling pathways (of 229) in the myometrium, which encompassed functions such as angiogenesis (HGF), homeostasis and tissue repair (CALCR, CLDN, PERIOSTIN, BMP), contractility (ncWNT), immune processes (CD70, IL10, FASLG, SEMA7, CD48, CD22, CSF3, CHEMERIN) and nervous system regulation (L1CAM, NGF) (**Supplementary Table 13**).

The loss of interactions driven by regulatory fibroblasts to endothelial cells through the HGF signaling pathway suggests impaired angiogenesis during aging (**Fig. 7A, Supplementary Table 13**).

**Figure 7.**
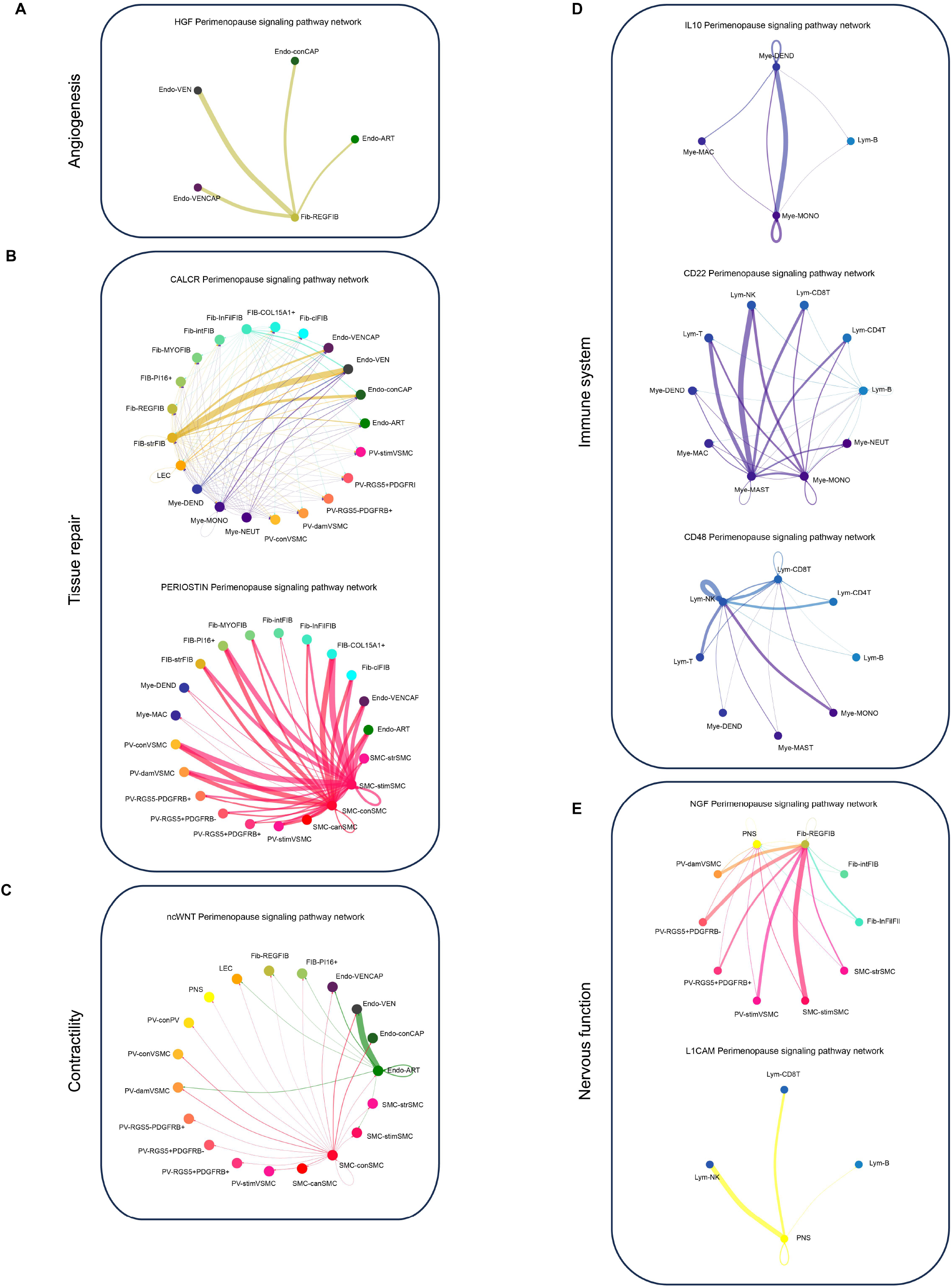
Lost Signaling Pathways in the Aging Myometrium. CCC chord diagrams displaying signaling pathways active in the peri-menopausal myometrium but lost during aging, including those related to **A)** angiogenesis (HGF), **B)** homeostasis and tissue repair (CLDN, CALCR, PERIOSTIN), **C)** contractility (ncWNT), **D)** immune processes (IL10, CD22, CD48) and **E)** nervous system regulation (NGF, L1CAM). Arrows between cell types show the direction of the interactions and follow the color code of cell types commonly detected in menopause. The relative thickness of each line depicts the expression-based strength of the interaction between cell types.

When evaluating pathways linked to tissue repair and homeostasis (**Fig. 7B**), we observed the loss of CALCR and PERIOSTIN signaling pathways in the post-menopausal myometrium - CALCR signaling functioned between myeloid cells and stressed fibroblasts in the peri-menopausal myometrium, while PERIOSTIN signaling was established by signals originating from stimuli and contractile SMCs. We also showed the loss of CLDN pathway in the post-menopausal myometrium (**Supplementary Table 13**), which was driven mainly by PI16+ fibroblasts in the peri-menopausal myometrium.

We also discovered the loss of non-canonical Wnt (nc-Wnt) during aging (**Fig. 7C**). The nc-Wnt signaling pathway primarily involved signals outgoing from contractile SMCs and arterial endothelium in the peri-menopausal myometrium. Importantly, nc-Wnt signaling in SMCs is essential for forming the actin-myosin-polarity structure required for contraction^26^.

When analyzing the impact of aging on immune system-related pathways (**Fig. 7D**), we discovered the loss of IL10 signaling pathway, (known for its potent anti-inflammatory properties) between myeloid cells and B-lymphocytes. Similarly, the CD22 pathway, which negatively regulates B-cell responses and involves signaling from mast cells, monocytes, and B-cells to other immune cells, was lost during myometrial aging. Lastly, we observed the loss of CD48 signaling (specifically the CD48-CD244 interaction), which normally plays a role in most immune cells and exhibits a strong autocrine signal in NK lymphocytes in the peri-menopausal myometrium.

Of note, we also observed the loss of pathways associated with nervous stimulation and transmission during myometrial aging, specifically through the nerve growth factor (NGF) and L1CAM pathways **(Fig. 7E).** The NGF pathway normally originates from specific types of PV cells, fibroblasts, stressed fibroblasts, as well as SMCs responsive to stimuli, and extended to the peripheral nervous system (PNS) and nervous system-regulatory fibroblasts **(Supplementary Fig. 4K)**, while the L1CAM pathway predominantly transmits signals from PNS to immune cells **(Supplementary Table 13).**

These findings highlight the complete loss of certain CCC in the aging myometrium, impacting multiple signaling pathways involved in angiogenesis, tissue repair, contractility, immune processes, and nervous system regulation.

Altogether our work provides a detailed atlas of the human myometrium and reports an age-related reduction in the proportion of contractile cells, and the impaired expression of genes in all cell types, such as voltage channels and contractility genes in SMCs and endothelial cells, ECM-related genes in fibroblasts, and inflammation-associated genes in immune cells. Further, myometrial aging disrupts interactions between critical cell types, resulting in defective angiogenesis, tissue repair, immunity, and nervous system regulation.

## Discussion

Aging represents a complex process encompassing altered tissue repair, fibrosis, tissue reprogramming, and cellular senescence^27^; unfortunately, reproductive organs are not exempt from adverse age-related changes. Our study which uses the progression of peri- into post-menopause as a model system, offers the first description of the interconnected cellular networks present in the human myometrium, providing evidence for altered CCC and the complete loss of crucial signaling pathways as hallmarks of myometrial aging. We document age-related endothelial dysfunction in the myometrium, which may contribute to microvasculature remodeling in conjunction with impaired angiogenesis and provide evidence for an age-related reduction in responsiveness to stimuli/electrical cues and disrupted signaling pathways involved in SMC contraction in the myometrium. Finally, the observed age-, induced fibrosis and perturbation to the immune system may prompt the compromised reproductive and obstetric functions observed in older women. We generated a comprehensive cellular landscape of the aging myometrium using single-cell and spatial transcriptomics, which revealed the transcriptomic activity of twenty-three specific myometrial subpopulations of cells. We discovered novel cell subpopulations, including contractile and venous capillary cells, characterized immune-modulated fibroblasts ^28, 29^, stressed fibroblasts, and nervous system regulatory fibroblasts^30, 31^, and identified stimuli-responsive SMCs, damage-and stimuli-responsive VSMCs, and immune cells. Overall, these findings highlighted the overall cellular heterogeneity of the aging myometrium.

When studying the endothelial cell population, we discovered an age-related reduction in the number of contractile capillary cells in the post-menopausal myometrium; however, we also observed altered function as revealed by the downregulated expression of contractility genes (such as *ACTG2* and *FLNA*) and the upregulated expression of genes linked to senescence and extracellular matrix deposition^32–34^. While we did not observe specific age-related changes in the abundance of fibroblasts, we did detect the upregulation of aging-associated genes (*FOS*, *FOSB,* and *NFATC2*) in universal PI16^+^ and COL15A1^+^ fibroblasts, classic fibroblasts, and intermediate NRP1+ fibroblasts during myometrial aging. SMCs also exhibited reduced functionality, as shown by the downregulated expression of contractility-associated genes (e.g., *ACTG2, MYL6, MYL9, TPM2,* or *DES*) despite an increase in their relative number, which may represent an attempt to compensate for the age-related loss of functionality and reduced contractility ^20, 32, 35^. Stimuli-response SMCs in the aging myometrium downregulated the expression of *KCNE4*, a type I beta subunit that assembles with the channel complex to modulate the gating kinetics and enhance stability^36^ and VDAC2, whose loss has been linked to cardiac muscle atrophy in mice^37^. The immune system plays a crucial role in senescence and aging, entailing impaired antigen presentation, T-cell function, and memory response leading to chronic inflammation (*inflammaging*) that disrupts organ functioning ^38, 39^. We observed the upregulated expression of genes related to immune cell activation, adhesion, migration, myelopoiesis, and monopoiesis (e.g., *PBX1, BNC2, EGR1, RORA,* and *DCN*) in the aging myometrium, suggesting disrupted immune homeostasis.

Our findings confirmed age-related alterations to CCC in the myometrium, which may prompt disturbances in interactions and physiological processes frequently observed as aging-related manifestations^40^. We demonstrated that significant differences in signaling pathways, the number of interactions, the specific ligand-receptor pairs involved, and a complete loss of several signaling pathways characterize the aging myometrium.

Of particular interest, we discovered the age-related loss of the HGF and NGF signaling pathways, which function in angiogenesis and blood flow^41, 42^. Such a loss could reduce HGF-mediated protective effects, which include wound-healing and anti-apoptotic/anti-inflammatory activities^43^; meanwhile, lost NGF signaling could impact physiological/pathological angiogenesis, through interactions with the vascular endothelial growth factor (VEGF)^44^.

Specific pathways that promote angiogenesis (e.g., ANGPTL, ANGPT, and VEGF^45, 46^) also displayed reduced activity during myometrial aging; we additionally discovered changes in the EDN pathway, which regulates blood flow through vessel constriction or relaxation. The predominant Endothelin Receptor Type A (EDNRA) ligand found in the post-menopausal myometrium has been linked to aging/aging-related diseases^47^ as demonstrated in other tissues and organs^48^. Of note, alterations to blood flow regulation and impaired neovascularization could impact the myometrium’s ability to control hemorrhage and nutrient delivery, potentially contributing to decreased functionality and increased morbidity and mortality in older women during pregnancy and labor^18, 49–51^.

The myometrium is mainly a muscular organ; therefore, reduced contractile force or increased stiffness caused by fibrosis significantly impacts normal function. Our study revealed a significant enrichment of fibrosis-related pathways during aging, which may compromise overall function. We observed increased PDGFC (implicated in fibrosis) and reduced PDGFD interactions, (associated with angiogenesis and blood vessel maturation)^52, 53^. The IGF pathway also exhibited subtle alterations, with stronger interactions between IGF1 and IGF1R (known for their role in pulmonary fibrosis^54^) in the post-menopausal myometrium.

We also detected changes in interacting ligand-receptor pairs of the NRXN signaling pathway, which helps to decrease sensitivity to stimuli interfering with impulse transmission and SMC contraction^55, 56^. The simplification/depletion of such CCC networks for various cell types in the aging myometrium may explain the decline in nerve impulse transmission and decreased contractility essential for pregnancy and labor physiology. Of note, some prior studies failed to encounter alterations in the contractile ability of myometrial tissues according to age^57^, while others have noted lower responsiveness to oxytocin stimulation^58^ or lower electromyographic activity^16^. Clinical studies have reported that older parturients have longer labors^59^, with higher dosages of oxytocin needed for third-stage labor to prevent uterine atony^60^. In this sense, our findings agree with decreased contractile potential during myometrial aging.

Our analyses revealed the absence of the ncWNT pathway in the postmenopause, which is involved in tissue homeostasis and cell migration^61^. The critical component WNT5A, known for its role in cardiac muscle contraction^62^, also strengthens actin cytoskeleton assembly^26^. Additionally, we observed a loss of CALCR signaling, which is crucial for maintaining muscle stem cells in a quiescent state^63^. Myometria unresponsive to stimuli or contractions can lead to uterine atony, resulting in postpartum hemorrhage^15, 64^ or labor dystocia.

Finally, we discovered that pathways involved in immune homeostasis, such as IL10^65^, CD22^66^ and CD48^67^ were absent in the post-menopausal myometrium. We also observed significant changes in the CXCL pathway - the post-menopausal myometrium displayed a decrease in CXCL2/8-ACKR1 interactions and an increase in CXCL12-CXCR4 interactions. While ACKR1 protects against inflammaging by limiting immune cell extravasation^68^, CXCR4 enhances the inflammatory response associated with aging-related DNA damage^69^. These changes suggest that impaired immune autoregulation in the aging myometrium may lead to a detrimentally overactive immune system. Inflammatory status conditions birth at term/preterm, as occurs with myometrial contractility orchestrated by cytokines released by immune cells that leads to parturition^70, 71^, while CXCL12 produced by the myometrium has been described as a causative factor for preterm labor^72^. Attenuating exacerbated myometrial immune responses may represent a therapeutic avenue to help reduce the incidence of preterm birth in older women^73^.

Of note, we acknowledge the limitations associated with our study. First, bias in cell/nuclei recovery could impact the likelihood of increased capture rates in coexisting populations, as observed in previous studies^74^. Second, the resolution of spatial transcriptomics data did not reach the level of individual cells or organelles, although we obtained comprehensive gene expression data for spots using unbiased high-throughput methods, which could hinder the visualization and estimation of minor cell populations. Nevertheless, integrating sc/snRNA-seq and spatial transcriptomics data is the most appropriate approach for uncovering diversity within myometrial cell populations and identifying spatial gene expression patterns.

Overall, our findings provide novel insight into the mechanisms underlying aging in the human myometrium; specifically, we highlight the diminished contractility of SMCs and endothelial and PV cells, disrupted immune responses, and changes in interactions within and between cell types, which impact inflammation, angiogenesis, fibrosis, and contractility. Ultimately, this knowledge may contribute to the development of preventive and therapeutic strategies to reduce pregnancy/labor complications in older women.

## Online content

Any methods, additional references, Nature Portfolio reporting summaries, source data, extended data, supplementary information, acknowledgments, peer review information; details of author contributions and competing interests; and statements of data availability are available at XXX.

## Methods

All experimental procedures and bioinformatic analyses in this work comply with ethical regulations and good scientific practices.

### Subject details

This prospective, multicenter, descriptive case series included twenty female donors, both living and deceased, with ages ranging from 46 to 79 years old. Our research employs the term “women” to refer to individuals with a uterus that undergo menopause while acknowledging that not all individuals identifying as women have a uterus and/or experience menopause, and not all individuals undergoing menopause identify as women. Patients were divided into two groups: the peri-menopausal group (46-54) and the post-menopausal (>54). In accordance with the definition of menopause as the absence of menstruation for a minimum of 12 consecutive months, our assumption, supported by prior studies^9^ and patients clinical reports, was that individuals aged 55 and above were post-menopausal. Conversely, patients aged 46 to 54 were categorized as peri-menopausal due to the likelihood of hormonal fluctuations and unpredictable menstrual cycle patterns during this phase. **Supplementary Table 1** describes the demographic and clinical characteristics of patients.

All procedures involving human tissue samples were approved by the Institutional Review Board of the Spanish hospitals involved: Hospital Clinico Universitario, Valencia, Spain (November 5th, 2019); Hospital La Fe, Valencia, Spain (December 4th, 2019); Hospital General Universitario, Valencia, Spain (February 12th, 2021). The Hospital La Fe provided fifteen uteri from women undergoing hysterectomy for pelvic prolapse, while a total of five uteri (two from Hospital Clinico Universitario and three from Hospital General) were obtained from deceased patients under the Organ Donor Program with non-cancer-related causes or traumatic injury.

All patients and donor families provided written informed consent, and those with gynecological disorders, malignancies, or diagnosed bacterial, fungal, or viral infections were excluded. This study was conducted per the International Conference on Harmonization Good Clinical Practice guidelines and the Declaration of Helsinki. All specimens were anonymized after collection and histologically evaluated by board-certified pathologists to confirm the diagnosis according to World Health Organization criteria.

### Tissue collection and sample preparation

Uterine tissues from the twenty recruited patients (n=3 samples per patient) were maintained in preservation solution (HypoThermosol® FRS) after surgery and further dissociated within 24 h of tissue retrieval to isolate the myometrial layer, which was achieved by removing the endometrium, serosa, and necrotic areas with a sterile scalpel. As previously described^75^, myometrial samples were rinsed in a wash buffer solution containing Hank’s Balanced Salt Solution (HBSS) (ThermoFisher Scientific, Grand Island, NY) and 1% antibiotic-antimycotic solution (ThermoFisher Scientific) to remove blood and mucus. Samples were then divided for different purposes; snap-frozen in liquid nitrogen or formalin fixation/paraffin embedding (FFPE) for further histological characterization, while the remaining tissue was carefully manually minced into small pieces (<1 mm^3^) and digested at 37°C using an enzymatic process for single-cell dissociation. Subsequently, cell suspensions were passed through a 50-mm polyethylene filter (Partec, Celltrics) to remove cell clumps and undigested tissue and then dissociated to single cells by incubating with 400μL TrypLE Select (ThermoFisher Scientific) for 20 min at 37°C to obtain single-cell suspensions. 100μL DNase I (Sigma-Aldrich) was then added to digest extracellular genomic DNA, and cells were treated with ACK Lysing Buffer (ThermoFisher Scientific) to induce hypotonic shock to avoid red blood cell contamination. The resulting cell suspension was pipetted, passed through a 100-μm cell filter, and centrifuged at 1000 rpm for 5 min. The cell pellet was resuspended in Dulbecco’s Modified Eagle’s Medium (DMEM, Sigma-Aldrich) supplemented with 2% fetal bovine serum (Labclinics) and 10 mM HEPES (ThermoFisher Scientific) as a myometrial-enriched suspension at a concentration of 1×10^6^ cells/mL. Cell concentration and viability were measured using trypan blue with an EVE™ automated cell counter (NanoEnTek). Dead cells were removed with the Dead Cell Removal Kit (Miltenyi Biotec) cells, with cell suspensions reaching a viability of ≥70%.

### Nuclei isolation for snRNA-seq from myometrial cell suspensions

A previously described snRNA-seq procedure^76^ was followed to increase the number of recovered cells from the human myometrium. After thawing and centrifuging cell samples, the supernatant was removed, and myometrial cell pellets were resuspended in 1 mL of lysis buffer on ice for 15 min. Samples were then transferred to a 15 mL conical tube and centrifuged at 500 g for 5 min at 4°C. The resulting cell pellets were resuspended in 1× ST buffer within the 100-200 µL range, and nuclei solutions were passed through a 40 µm Falcon cell strainer. The concentration and viability of nuclei were evaluated using an EVE™ automated cell counter (NanoEnTek) with trypan blue. Finally, ∼10,000 single-nuclei suspensions were loaded onto Chromium Chips for the Chromium Single Cell 3′ Library preparation according to the manufacturer’s recommendations (10x Genomics).

### Single-cell capture, library preparation, and sequencing

To profile single cells/nuclei from three different areas of the human myometrium (anterior, posterior, and fundus), scRNA-seq analysis was performed using the 10X Chromium system (10X Genomics, Pleasanton, CA, USA). Approximately 17,000 cells or nuclei were loaded onto a 10X G Chip to obtain Gel Bead-in-emulsions (GEMs) each containing an individual cell. GEMs were used to generate barcoded cDNA libraries following the manufacturer’s protocol (Single Cell 3’ Reagent Kit v3.1, 10X Genomics) and quantified using the TapeStation High Sensitivity D5000 kit (Agilent, Germany). Subsequently, gene expression libraries were constructed using 1-100 ng of each amplified cDNA library and quantified using the TapeStation High Sensitivity D1000 kit (Agilent, Germany) to determine the average fragment size and library concentration. Libraries were normalized, diluted, and sequenced on the Illumina NovaSeq 6000 system (Illumina, USA) according to the manufacturer’s instructions.

### Spatial transcriptomics

The systematic mapping of cells and gene activity to tissue locations used the Visium Spatial Gene Expression Reagent Kit (10X Genomics). Eight full-thickness uterine samples were examined from eight women - three peri-menopausal (under 55 years old) and five post-menopausal (equal to or older than 55 years old).

First, the quality of preserved RNA in FFPE blocks was evaluated based on the percentage of RNA fragments above 200 base pairs (DV200). Next, 7 µm-thick tissue sections were sampled using a semi-automated microtome (ThermoScientific HM340E). Per the manufacturer’s protocol, each section was mounted onto a 6.5×6.5 mm capture area of the Visium Spatial Gene Expression Slide (PN-2000233). Capture areas contain approximately 5,000 barcoded spots, providing an average resolution of 1-10 cells (10X Genomics).

Tissue sections were then deparaffinized, hH&E stained, and uncrosslinked according to the manufacturer’s protocol (10X Genomics) with minor modifications. Brightfield images were taken using a 10X objective (Plan APO) on a Nikon Eclipse Ti2. Images were stitched together using NIS-Elements software (Nikon) and exported as .tiff files. Following imaging and uncrosslinking, the Visium Spatial Gene Expression Slide & Reagent for FFPE kit (10XGenomics) was used for library construction. All steps (cDNA synthesis and amplification, library construction, and post-library construction quality control) were carried out according to the manufacturer’s protocol. The libraries were sequenced on a HiSeqX (Illumina), 50PE (2x 150bp), applying 1% Phix. Sequencing depth was calculated with the formula (Coverage Area x total spots on the Capture Area) x 50,000 read pairs/spot. Sequencing was performed using the following read protocol: read 1: 28 cycles; i7 index read: 10 cycles; i5 index read: 10 cycles; read 2: 90 cycles. Following sequencing, data was visualized to determine each gene expression’s spatial location and degree.

### sc/snRNA-seq data processing and filtering

Raw sequences were demultiplexed, aligned, and counted using the CellRanger software suite (v 6.0.2) for nuclei and whole cell gene expression calculations, which takes advantage of intronic reads to improve sensitivity and sequencing depth (human reference genome GRCh38-2020-A). Technical artifacts due to ambient RNA contamination were reduced with CellBender (0.2.0), and low-quality droplets and barcodes were filtered out in four quality control-based consecutive steps throughout the analysis: (i) low UMI-count barcode removal using an EmptyDrops-based method; (ii) cells/nuclei marked as doublets by DoubletFinder (2.0.3) and scds (1.6.0) tools – the hybrid approach from the scds R package was used to avoid removing false-positive doublets; (iii) cells/nuclei with median absolute deviation (MAD) > 3 in two of three basic quality control metrics: number of detected features, number of counts and mitochondrial ratio. These cell-to-count matrices were integrated and corrected using Seurat and Harmony functions, as described below. A final filtering step, (iv), was applied alongside different rounds of clustering, where the obtained clustered cells/nuclei with less than 750 features/cell, more than 25% mitochondrial ratio, and/or showing a pattern of high doublet-scoring plus no gene marker associated expression (during manual cell type annotations), were also removed.

Nuclei droplets were similarly processed, and quality controls were applied with the only specific criteria to remove nuclei with MAD > 2 in mitochondrial ratio per sample.

### Integration of single cells and single nuclei across conditions and clustering

As a first clustering analysis approach, read count matrices per sample were merged and processed following Seurat’s default pipeline (package version 4.1.3). After normalization, the first thirty principal components on the 4,000 highly variable genes were used for dimensional reduction; cells were clustered and projected onto the UMAP. *FindNeighbors* and *FindClusters* functions were then applied for graph-based clustering by constructing a KNN graph using Euclidean distance in the principal component analysis space, which was then defined into clusters using the Louvain algorithm to optimize the standard modularity function. clustree (R package v0.4.4) was applied to select the most stable clustering resolution. The first output of sample distribution in clusters and cluster marker genes was then explored to evaluate biases from our data batches. Next, the Harmony R package (v1.0) was used to remove four primary sources of bias: i) between single-cell and single-nuclei protocols, ii) between tissue zone of origin (anterior, posterior, or fundus), iii) between aging conditions, and iv) the patient sample origin to remove inherent inter-individual differences. Dimensions were reduced as before, and an integration diversity penalty parameter (theta) of 2 was used. The top thirty Harmony components were used to embed and plot cells in the new reduced dimension space. These matrices were then input into Seurat’s clustering and differential expression protocol. The clustering of different primary cell types for fine-grain cell type annotations was equally computed, following all the above-described steps.

### Tissue cell composition analysis and annotation of sc/snRNA-seq datasets

The identification and labeling of major cell types used the analysis of differentially expressed genes of each cluster compared to the remaining clusters using the Wilcoxon Rank Sum test and adjusted p-values for multiple comparisons with the false discovery rate (FDR) method^77^. The primary cell populations were labeled by revising the expression of reported canonical markers from each cluster. The five main cell types were then subset separately: endothelial cells, fibroblasts, SMCs, PV cells, and immune cells, and a new clustering was performed on each to create the different zoom-ins that describe the contained sub-populations. The clustered zoom-ins were manually annotated by an extensive review of differentially expressed genes in the Human Protein Atlas database, single-cell atlases, and scientific literature^23, 74, 78–86^. Genes with strong cluster-specificity, as determined by a p-value below 0.01, and the highest rank fold change and percentage of expressing cells were considered. Over-representation analysis on functional gene ontology and Reactome terms was also performed on cluster-specific genes with the WebGestaltR package (version 0.4.5) to help interpret cluster biological functions.

### Analysis of differential cell abundances and gene expression between peri and post-menopausal myometria

Identifying cell subpopulations with differential abundance comparing peri-menopausal and post-menopausal myometria as a model of aging used the Milo approach described by Dann et al.^87^, which is available as a miloR package (version 1.6.0). This approach supports the grouping of cells on a k-nearest neighbor graph and evaluates the change in cell abundance between peri-menopausal and post-menopausal conditions. Further differential gene expression analysis between conditions was performed using the Model-based Analysis of Single Cell Transcriptomics (MAST), where a contrast test was established for each cell type^88^. The over-representation analysis on aging-associated genes identified enriched gene ontology biological processes and Reactome pathways.

### Spatial transcriptomics data processing

Spatial transcriptomics data processing used the Space Ranger v2.0.1 to align sequence data to the GRCh38-2020-A reference genome, followed by tissue and fiducial detection of spots and counting barcodes/unique molecular identifiers (UMIs) to generate feature-barcode matrices. These matrices were loaded into Seurat, and quality control filtering was performed based on the number of reads and detected genes per spot. Next, a reference matrix was built with informative genes from scRNA-Seq data (with a mean expression in a specific cell type at least 0.75 log fold higher than other cell types and removal of top 1% genes with the highest expression dispersion). Conditional autoregressive-based deconvolution was used to estimate cell composition in each spatial transcriptomics spot and build refined high-resolution maps of cell proportions and gene expression using the CARD package^89^. Cell proportions estimated by CARD were normalized using the min-max method to validate differential abundance detected by scRNA-seq, while the spatial count matrix was processed using total count and log-normalization for validating differential expression detected by scRNA-seq. These data were then plotted in boxplots and contrasted differences between peri-menopausal and post-menopausal myometrial samples by a Wilcoxon signed-rank test.

### Analysis of Cell-Cell Communication

Identifying potential cell interactions between different cell populations in the aging human myometrium used the CellChat R package (v1.1.3)^22^. This tool infers the total interaction probability and communication information flows based on the expression of specific ligand-receptor pairs supported by a curated database. Briefly, the total interaction probability represents the probability of communication between two cell types, one acting as the sender and the other as the receiver, based on the number of interactor molecules expressed (ligand-receptor pairs) and the strength of this interaction (expression). Then, the sum of the communication probability of all pairs in a pathway network is used to calculate the communication information flow. For this analysis, cells labeled as cycling and pathways with less than ten cells were removed, and the influence of each cell population size was corrected by setting the *population.size* argument in the *computeCommunProb* function to *TRUE.* Lastly, differential CCC analysis between peri- and post-menopausal myometria was performed with the *ranknet* function and a significance threshold of 0.05.

## Material availability

This study did not generate new unique reagents. Material that can be shared will be released via a Material Transfer Agreement. Further information and requests for resources and reagents should be directed to and will be fulfilled by the lead contact Aymara Mas at.

## Reporting summary

Further information on research design is available in the Nature Portfolio Reporting Summary linked to this article.

## Data Availability

Sequencing data supporting this study’s findings have been deposited in the Gene Expression Omnibus (GEO). Source data are provided with this paper. All other data supporting the findings of this study are available from the corresponding author upon reasonable request.

## Acknowledgments

This study was jointly supported by the H2020-funded project Human Uterus Cell Atlas (HUTER 2020/2021) (Grant Agreement 874867), Miguel Servet Spanish Program Grant (CP19/00162), Health Research Funds (PI20/00942) from Carlos III Institute, Spain (AM), as well as Generalitat Valenciana (FDEGENT/2019/010) and (ACIF/2021/348) Ph.D. Training Grant for Valencian Entities (AML/PPJ). R.P. was supported by an Industrial Doctorate grant (DIN2020-011069) from the Spanish Ministry of Science and Innovation (MICINN). We acknowledge Javier Monleón, Ana Ochando, Stuart P. Atkinson, and Carlos Lozano for their contributions to this study.

## Author Contributions

Conceptualization: AM, CS. Formal analysis: AML, PPJ, JL, RP, BR. Data curation: AML, JL, RP, BR, EP. Investigation: AML, PPJ, MG. Resources: ES, CF, RB, DG. Validation: AML, MG, PPJ, RP, BR. Writing – Original Draft: AML, PPJ. Writing – Review & Editing: AM, CS. Supervision: AM, CS. Funding acquisition: AM, CS. All authors have substantively revised the manuscript and approved the submitted version.

## Additional information

**Extended data** is available for this paper at XXX.

**Supplementary information** is available for this paper at XXX.

**Correspondence and requests for materials** should be addressed to AM.

## Supplementary Figures

**Supplementary Figure 1.**
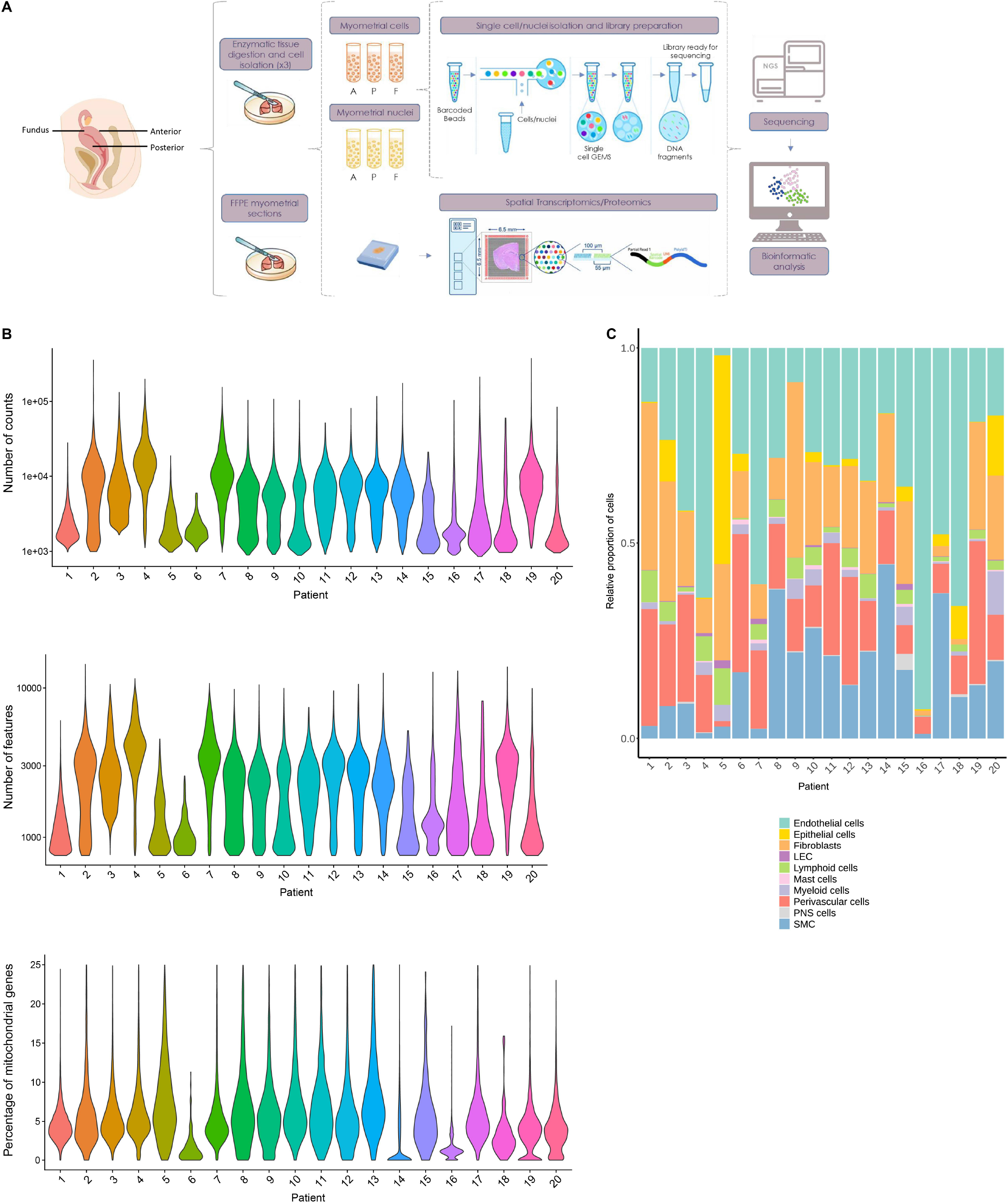
Single-cell Profiling of the Human Myometrium. **A)** Overview of the study workflow. **B)** Violin plots illustrating the quality control metrics used in the sc/snRNA-seq analysis for each patient’s features, counts, and mitochondrial percentage per cell. **C)** Bar plots illustrate the relative contributions of myometrial cell types in each patient. Abbreviations: LEC: lymphatic endothelium; PNS: peripheral nervous system; SMC: smooth muscle cell.

**Supplementary Figure 2.**
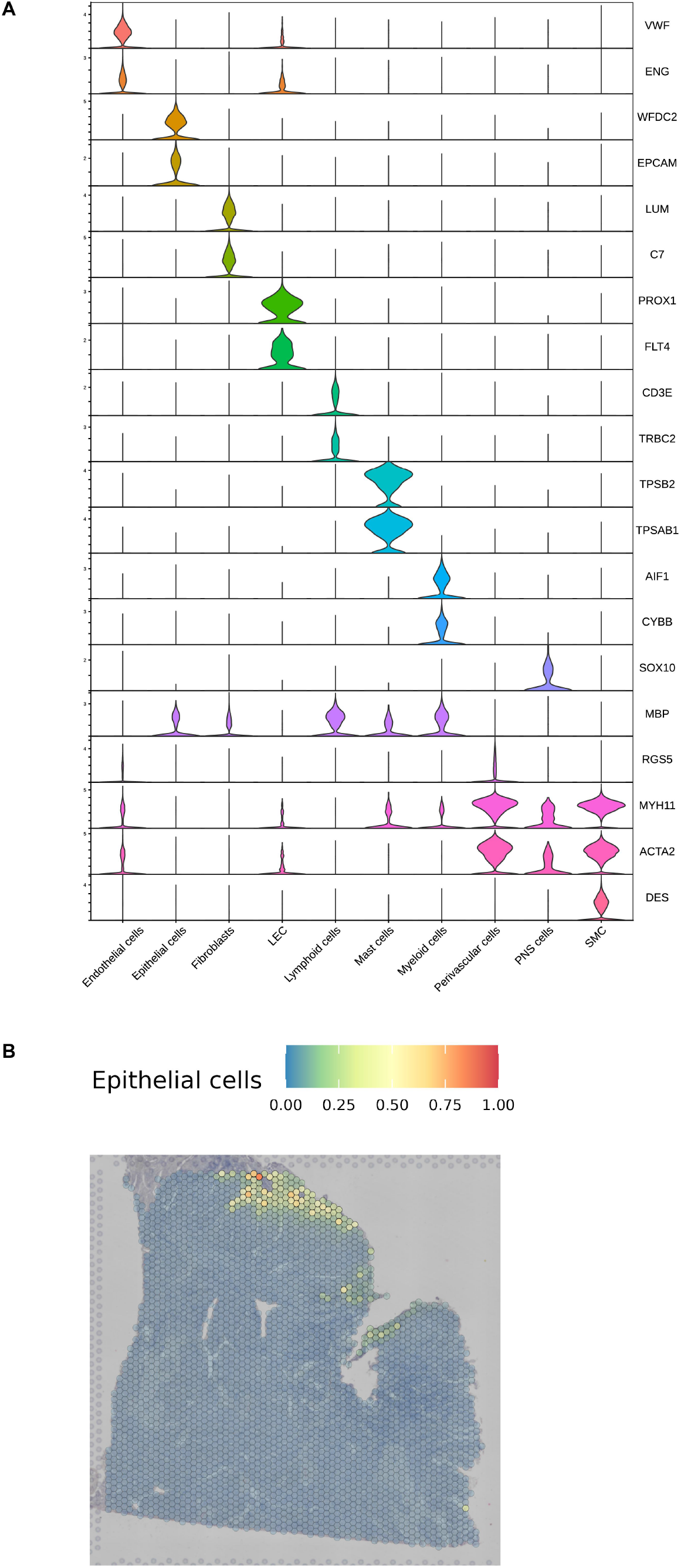
Single-cell Expression of Canonical Marker Genes and the Spatial Localization of Epithelial Cells within the Myometrium. **A)** Violin plots reporting the expression of canonical marker genes for main cell types. **B)** Representative image of the spatial distribution of epithelial cells within the myometrial tissue. Spot color indicates the proportion of epithelial cells in each spot.

**Supplementary Figure 3.**
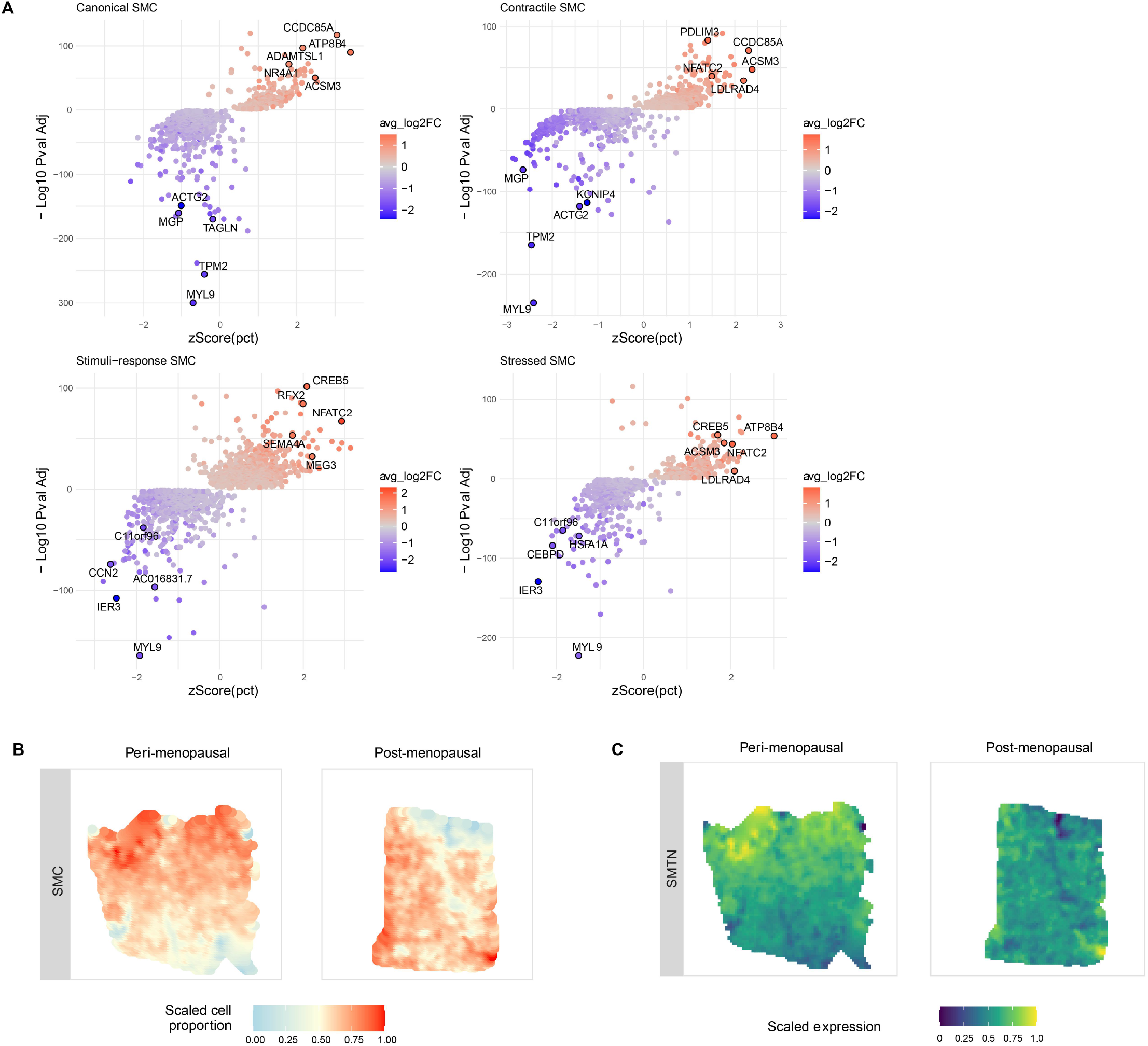
Comparison of Smooth Muscle Cell Subtypes in Peri- and Post-menopausal Myometrial Samples. **A)** Volcano plots representing differentially expressed genes in the four SMC subpopulations during myometrial aging. **B)** Representative refined spatial maps of SMCs in peri-(left) and post-menopausal (right) myometrium. **C)** *SMTN* gene expression integrated onto a refined spatial map of the peri- and post-menopausal myometrium.

**Supplementary Figure 4.**
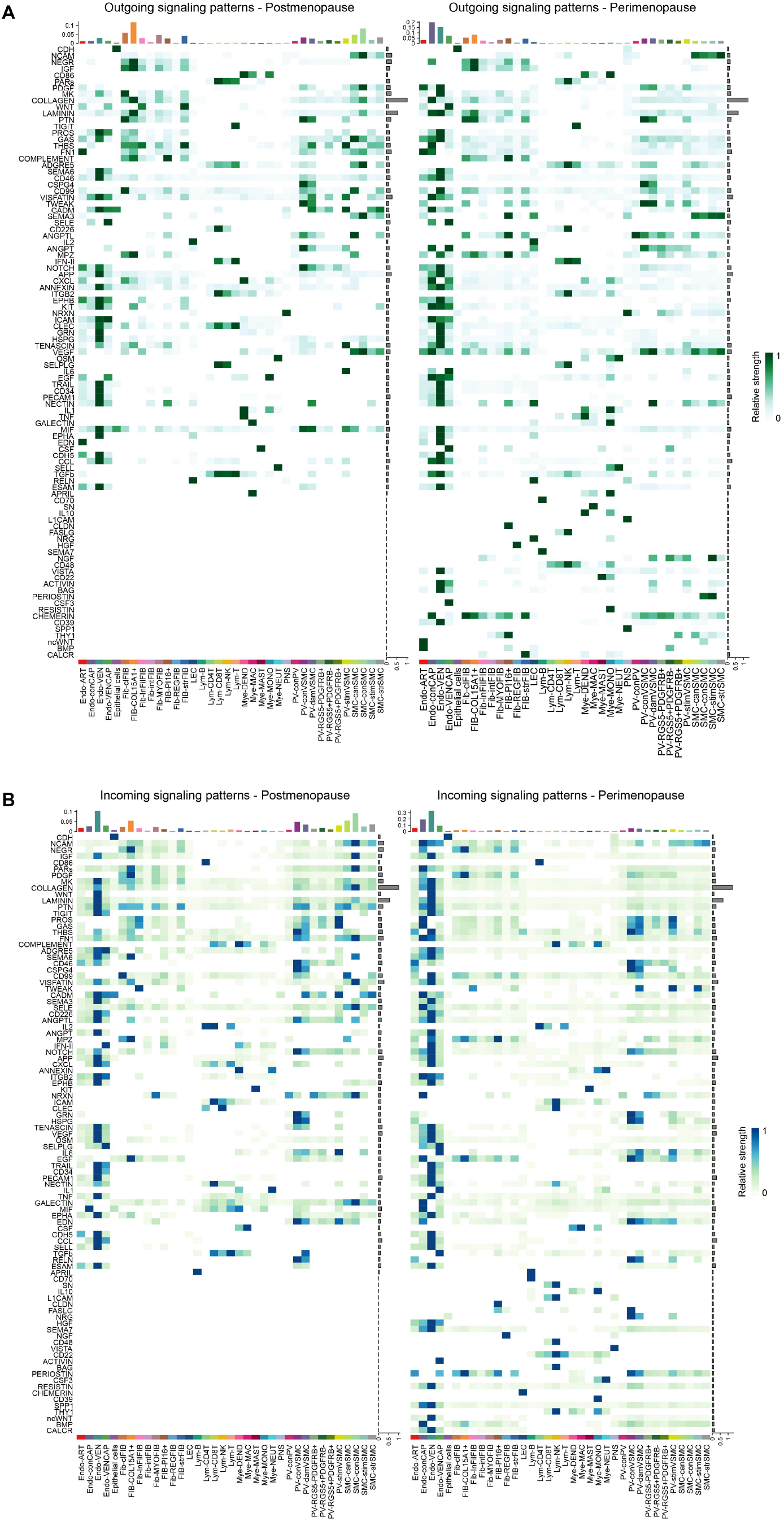
Differential Age-Related Changes in Cell-to-Cell Communication in the Myometrium. **A and B)** Heatmaps report the contribution of signaling pathways in each cell subtype in terms of **A)** outgoing signaling and **B)** incoming signaling in the peri- (left) and post-menopausal (right) myometrium. The y-axis reports the signaling pathway, while the x-axis represents the number of cells of each subtype to which the relative strength of the signal refers (top) and the subtype name displayed (bottom). The colored squares report the relative strength of each signaling pathway.

## Supplementary Tables

**Supplementary Table 1.** Epidemiological, demographic, and clinicopathological outcomes of peri-menopausal (n=6) and post-menopausal (n=14) women included in the study

**Supplementary Table 2.** Run summary metrics of single-cell libraries in the menopausal human myometrium.

**Supplementary Table 3.** Descriptive statistics of marker genes associated with endothelial subclusters.

**Supplementary Table 4.** Differentially expressed genes in endothelial subclusters during myometrial aging (Adjusted p-value<0.05; avg_log2FC>0 indicates overexpression in post; avg_log2FC<0 indicates downregulation in post-menopause).

**Supplementary Table 5.** Descriptive statistics of marker genes associated with fibroblast subclusters.

**Supplementary Table 6.** Differentially expressed genes between fibroblasts subclusters during myometrial aging (Adjusted p-value<0.05; avg_log2FC>0 indicates overexpression in post-menopause; avg_log2FC<0 indicates downregulation in post-menopause).

**Supplementary Table 7.** Descriptive statistics of marker genes associated with SMC subclusters.

**Supplementary Table 8.** Differentially expressed genes between smooth muscle subclusters during myometrial aging (Adjusted p-value<0.05; avg_log2FC>0 indicates overexpression in post-meonpause; avg_log2FC<0 indicates downregulation in post-menopause).

**Supplementary Table 9.** Descriptive statistics of marker genes associated with PV subclusters.

**Supplementary Table 10.** Differentially expressed genes between PV subclusters during myometrial aging (Adjusted p-value<0.05; avg_log2FC>0 indicates overexpression in post-meonpause; avg_log2FC<0 indicates downregulation in post-menopause).

**Supplementary Table 11.** Descriptive statistics of marker genes associated with immune subclusters.

**Supplementary Table 12.** Differentially expressed genes between immune subclusters during myometrial aging (Adjusted p-value<0.05; avg_log2FC>0 indicates overexpression in post-meonpause; avg_log2FC<0 indicates downregulation in post-menopause).

**Supplementary Table 13.** Relative contributions of ligands and receptors to signaling pathways during myometrial aging. The first column reports the pathway name, the second column the ligand-receptor pairs contributing to the signaling of each pathway, and third column displays the values for relative contributions in terms of percentage that each pair ligand-receptor has within the pathway.

